# scMetric: An R package of metric learning and visualization for single-cell RNA-seq data

**DOI:** 10.1101/456814

**Authors:** Wenchang Chen, Xuegong Zhang

## Abstract

Single cell RNA-seq data provide high-dimensional transcriptome features of each cell and enables the systematic study of cell types and cell states based on molecular characteristics, which are usually regarded as clusters in the high-dimensional space. Usually not all genes are equally informative for the studied biological question and the clusters are embedded in some lower dimensional subspaces.

There are several popular dimensionality reduction and visualization methods for scRNA-seq data such as PCA and t-SNE^1^. Different methods may utilize different kinds of distance metrics. Some methods take distance metric as a hyper-parameter for users to choose. Different metrics can result in different data distribution in the low dimensional space^2^ and therefore result in different final results. But there is no way to know which metric is “the correct one” for the particular biological study at hand. A typical practice is to use the default metric provided by the software and wait to see whether results look compatible with existing knowledge or users’ expectation. Gene selection steps are usually adapted before applying the metric to filter those genes that are less likely to be relevant to the study, and/or use supervised information (e.g., the classification of cell types based on known markers) to select genes. Such strong supervised information may cause the results to be overly dependent on the existing knowledge, which may then reduce the possibility of new discoveries.

In studies that aim to profile the composition of cell types and cell states of a complicated physiological or pathological system, there could be multifaceted underlying relations among the cells according to different angles of study. For example, cells can be clustered by their development stages and also by their lineages in developmental studies^3^. A fixed metric for dimensionality reduction and other downstream analyses does not provide the flexibility of exploring the data from multiple angles.

We proposed a machine learning method to allow users to only provide weak supervision and let the machine to learn a proper metric to be used on the genes, based on the idea of metric learning^4,5^. The weak supervision can be as simple as a few example pairs of similar cells and of dissimilar cells. In this way, users only use a small portion of confident examples to train the method, without introducing pre-assumptions on most of the cells. We developed the package scMetric (Fig. 1) to apply metric learning on gene expression data. The package let users to assign a few pairs of cells that are similar with each other and another few of dissimilar cells. With this weak training, scMetric learns the metric A to best preserve the similarity and dissimilarity reflected in the training pairs. It then employs t-SNE to visualize the data using this metric. The package outputs the t-SNE maps using the learned metric and using the conventional Euclidean distance metric. The learned metric A as well as the key genes that compose most weights in the metric can also be output to other analysis methods that need a distance or similarity metric.

**Fig. 1.**
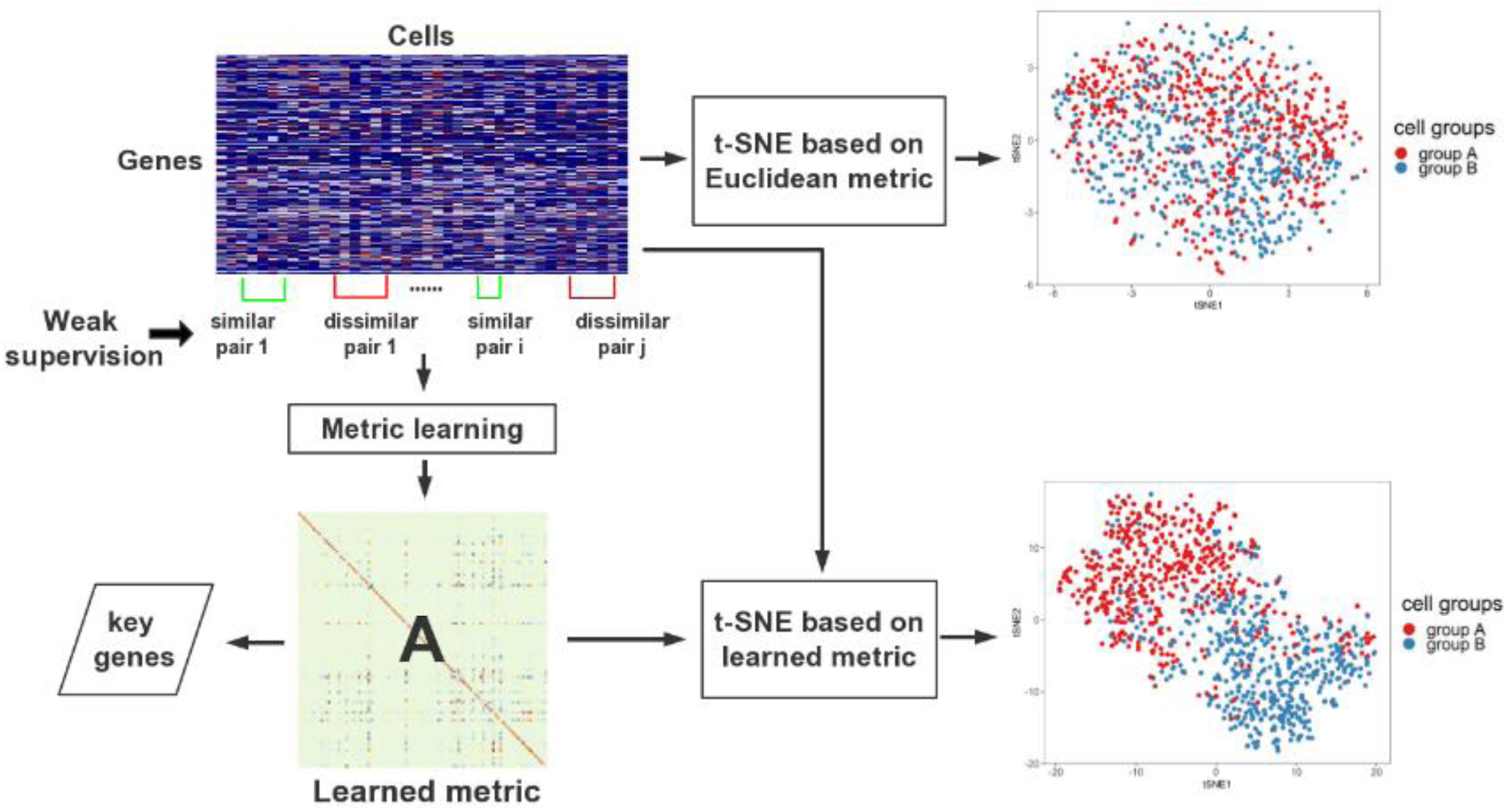
Single cell RNA-seq data metric learning workflow. Inputs are a gene expression matrix as well as the weak supervision information. The t-SNE maps shows examples of the visualization results after metric learning and without using the learned metric.

We did simulation and real-data experiments to show advantage of the method. In the simulation, we seeded two orthogonal schemes of classification: Cells are categorized to classes A or B with some genes. With another set of genes, the cells are classified to classes C or D. Each cell has two class labels and they are not correlated.

Noisy genes are added so that either the distinction between A and B or the that between C and D are dominate in the data, and the direct visualization of t-SNE based on the Euclidean metric cannot reveal either classes. Using scMetric, if the training pairs are chosen according to the grouping of A and B, the t-SNE map with the learned metric separates classes A and B well. On the other hand, if we choose training pairs according to the distinction of C and D, then these two clusters become clear in the t-SNE map. We applied the method on a real dataset of 6 clusters and two experiment batches. Results show that it can effectively capture different aspects of the major information based on different weak supervision.

Details of the experiments as well as the method are described in the Supplementary File. The R code of scMetric is available on GitHub at https://github.com/XuegongLab/scMetric. The package can also be used for other types of data that have similar tasks.

## Author contributions

Wenchang Chen wrote scMetric and analyzed data; Xuegong Zhang planned the study and wrote the manuscript.

## Acknowledgements

This work is supported by CZI HCA pilot project, the National Key R&D Program of China grant 2018YFC0910400 and the NSFC grant 61721003.

## Additional Information

Supplementary information on the method and experiments are attached.

## Supplementary information

### Methods

ITML (Information-Theoretic Metric Learning, Davis, *et al*. 2007) is a classic metric learning method based on Mahalanobis distance. It solves a Bregman optimization problem that minimizes the differential relative entropy between two multivariate Gaussians under distance constraints. The goal is to learn the positive definite A in distance function
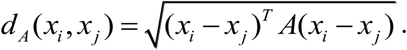

Usually, parameter matrix A is regularized to be as close as possible to a known matrix A_0_ like identity matrix or Mahalanobis matrix. If we assume data obeys Gaussian distribution, it may be better to choose Mahalanobis matrix. In other cases, identity matrix may have better performance according to experiments.

### From Matric distance to Kullback–Leibler divergence

Then the next problem is that how to measure similarity between two matrices A and A_0_ so that we can ensure A is closer to A_0_ step by step during the learning. From statistic knowledge we know that there exists a bijection between the set of equal mean multivariate Gaussian distribution and the set of Mahalanobis distance. Therefore, we can minimize distance between two multivariate Gaussian distribution to represent distance between two matrices. The corresponding multivariate Gaussian for matrix A is
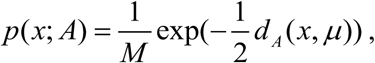

where *μ* is the mean, and M is a normalization factor. KL divergence is widely used to measure distance between two distributions. The distance between two multivariate Gaussian for matrix A and A_0_ is
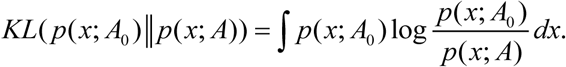

#### From Kullback–Leibler divergence to LogDet Optimization

The LogDet divergence for *n* × *n* matrices A and A_0_ is a Bregman matrix divergence that equals to
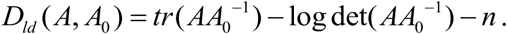

Davis and Dhillon (2006) showed that KL divergence of two distributions can be expressed as the convex combination of a Mahalanobis distance between their mean vectors and the LogDet divergence between their covariance matrics. Assuming that two Gaussians have the same mean, KL divergence can be expressed as
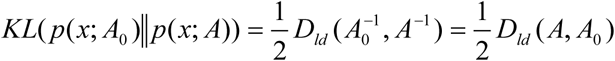

#### Constraints

Suppose we have a dataset that consists of n samples {*x_1_, x_2_, …, x_n_*} and two sets of sample pairs of S and D. If a pair of samples are similar according to prior information, we put it in S. Otherwise we put it in D.
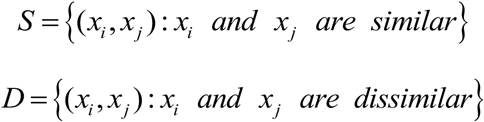

Given S and D, two values *u* and *l* should be decided. *u* is a relatively small value that represents upper distance bound of pairs in S. *l* is a relatively large value that represents lower distance bound of pairs in D. To estimate a proper choice of *u* and *l*, we can calculate the distances between all possible and non-repeating cell pairs. Usually, half of them, *n*\*(*n*-1)/2 pairs, are selected randomly in consideration of speed. In case of *n*\*(*n*-1)/2 is still too large, we set an upper bound of 10,000. Then we calculate distance of these selected pairs and assign 5% and 95% of the largest distance to *u* and *l*, respectively.

The constraints are set as the following:
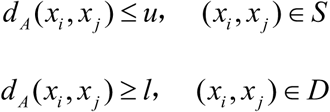

The final optimization problem is:
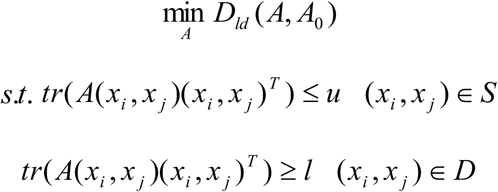

#### Finding key genes from learned new metric A

New metrics can be seen as transformations of space that assign weights on genes differently. Since A is a nonnegative positive definite matrix, it can be decomposed as
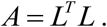

Then new distance is
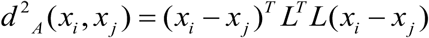

Let
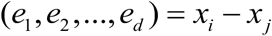

Assume that only the *k*-th gene expression changes while others remain the same
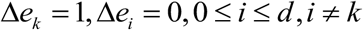

This results in change of distance
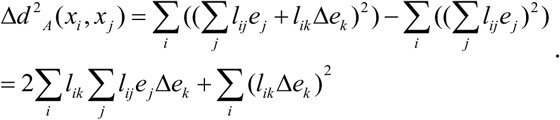

We compare |Δ*d*^2^*_A_*(*x_i_, x*_j_)| for different genes. The larger it is, the more weights new metric assigns on genes. We use this measurement as the importance score to order the relative importance of genes in the learned metric.

## Simulation Experiments

We first applied our method to a group of simulated data. Figure S1 is an illustration of the main idea of the simulation experiment. Assume that there exist two kinds of grouping criteria in the simulated data. We can separate cells into group A and group B according to criteria 1 while criteria 1 will not be able to separate group C and group D. On the other hand, criteria 2 separates group C and D without considering group A and B.

**Fig. S1.**
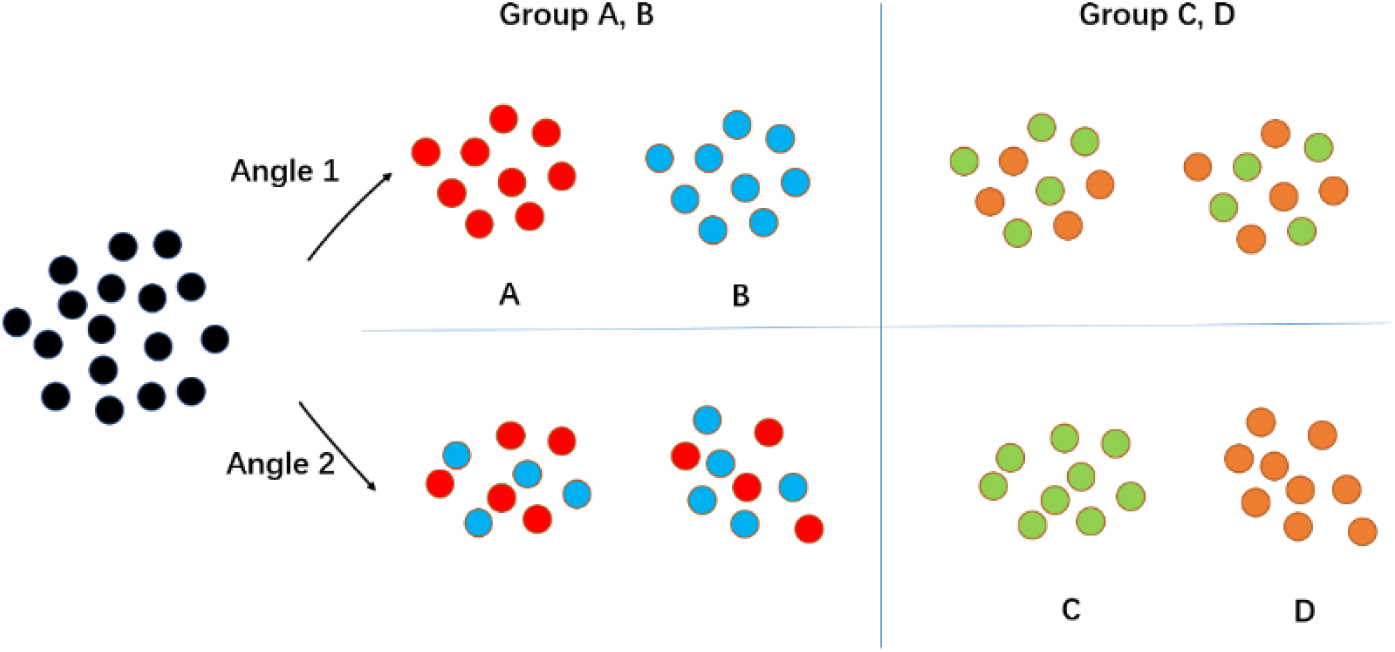
Illustration of the simulation data design.

We used *splatter* (Zappia et al, 2017) to simulate single cell data. We simulated 1000 cells with 1000 genes selected. Among them, 100 genes have 30% probability to differentially expressed between group A and B, and another 100 genes have 30% probability to differentially expressed in group C and D. The rest 800 genes are randomly simulated as noisy genes. Figure S2 shows t-SNE maps of simulated cells with different genes.

**Fig. S2.**
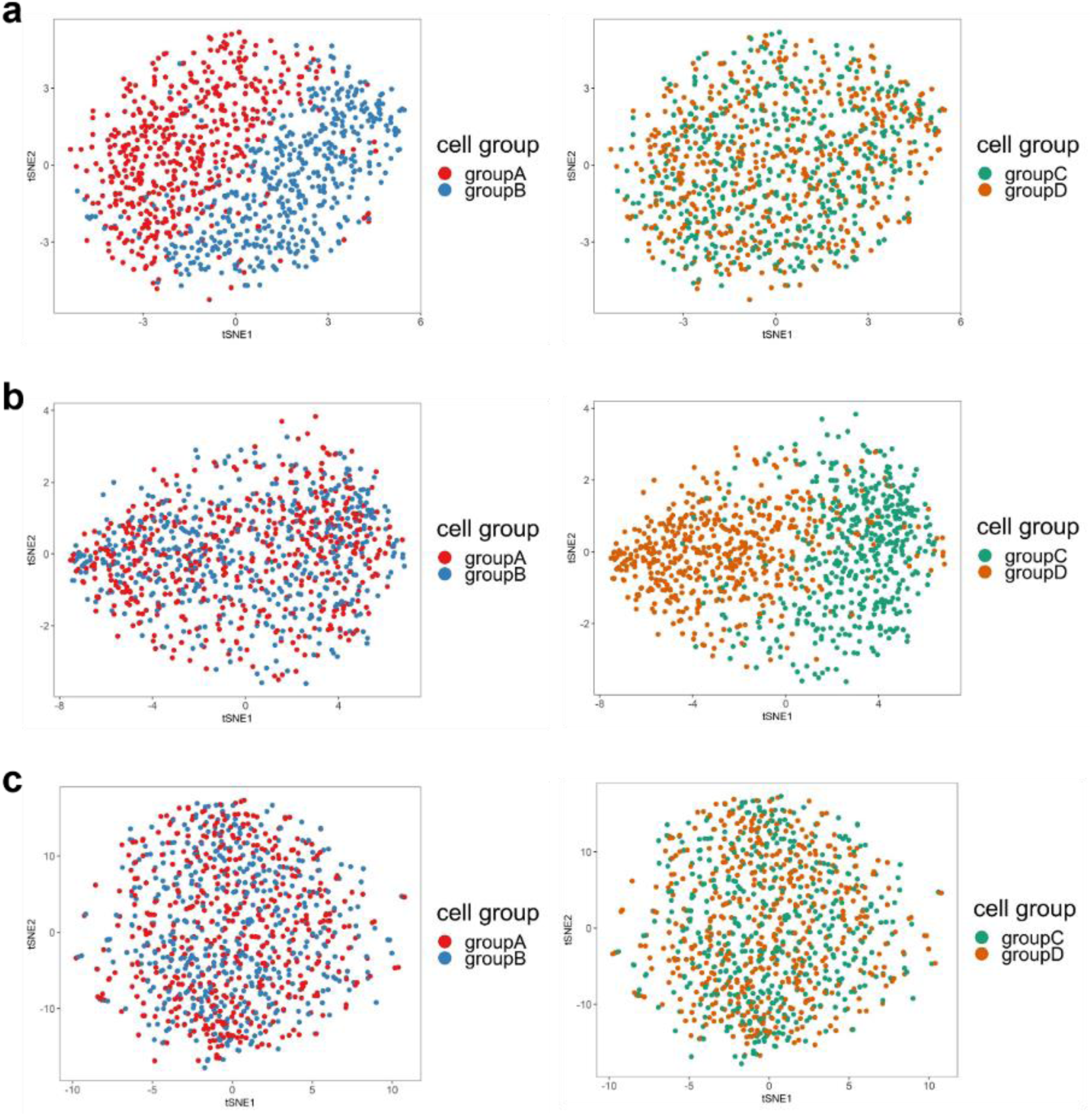
t-SNE maps of 1000 simulated cells with different gene sets. The two t-SNE maps of each row are the same but the cells are colored by groups A and B in the left panel and groups C and D in the right panel. (a) t-SNE maps with 100 genes that contain DE genes between groups A and B. (b) t-SNE maps with 100 genes that contain DE genes between groups C and D. (c) t-SNE maps with 800 noisy genes.

Figure S3(a) shows t-SNE maps of 1000 cells with all 1000 genes. We can see that since the DE genes between any of the two groups are only a small portion of all genes, the signals are weak and the direct visualization of t-SNE cannot reveal any classes. Then we choose 50 training pairs (25 similar pairs and 25 dissimilar pairs) for each grouping scheme and apply metric learning on original data. Figure S3(b) shows t-SNE maps scMetric outputs based on learned metric rather than conventional Euclidean distance metric. We can see that the t-SNE maps can well separate the corresponding two groups of cells according to the weak supervision information, without involving any gene selection procedure.

**Fig. S3.**
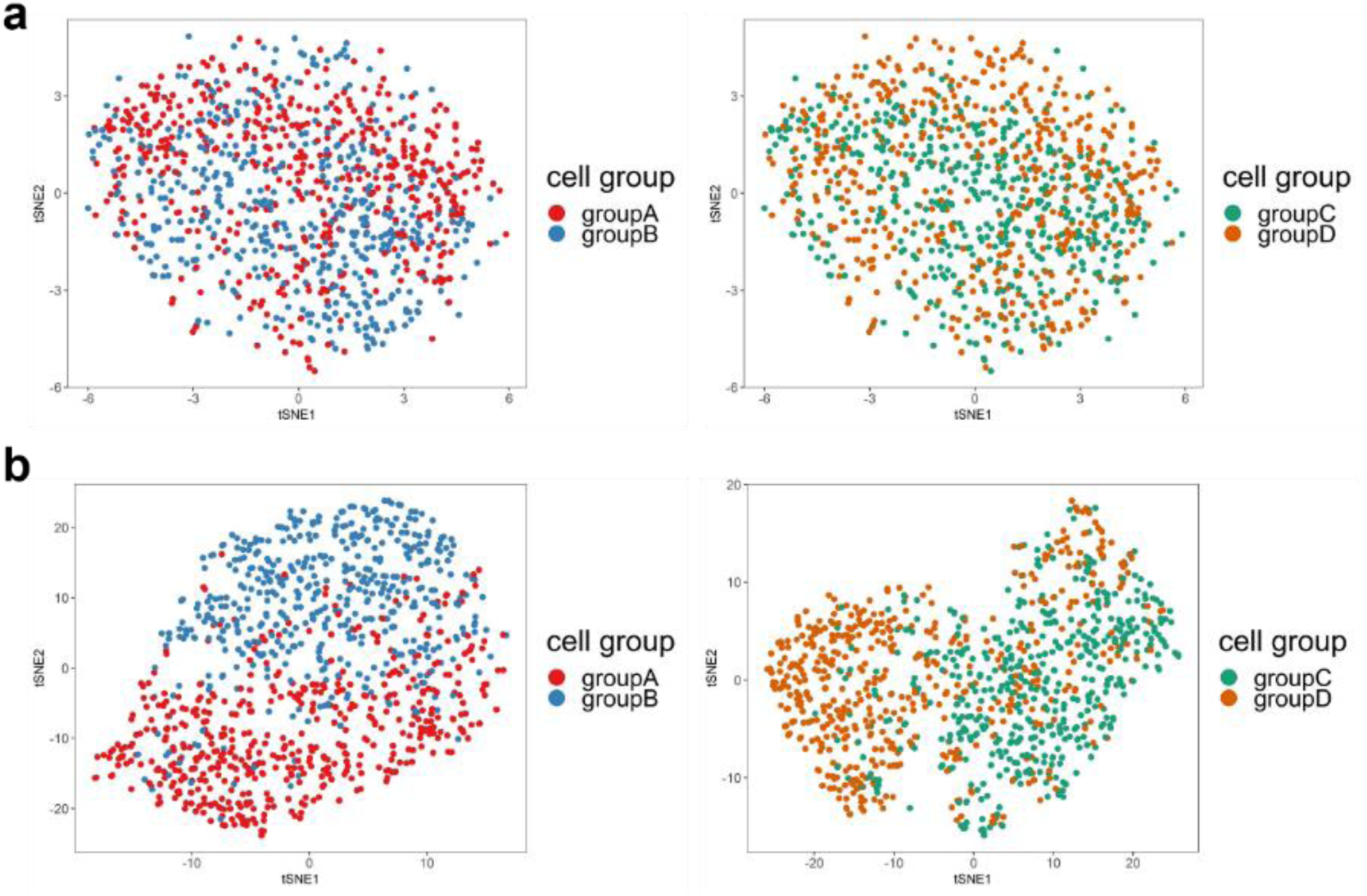
t-SNE maps shows the visualization with and without using the learned metric by scMetric. (a) t-SNE maps with Euclidean metric. (b) t-SNE maps with the learned metric using weak supervision for groups A and B (left) and with learned metric using weak supervision for groups C and D (right).

## Real Data Experiments

### Single-cell RNA-seq data of dendritic cells (DCs)

We applied scMetric on a single-cell RNA-seq dataset of dendritic cells (DCs) to study its effectiveness on real data. In Villani et al. (2017), the authors identified 6 subtypes of dendritic cells and 4 monocyte populations through single cell RNA-seq data. We did the experiment on the 742 dendritic cells in their dataset. After gene filtering following the preprocessing procedures in the original paper, we obtained a gene expression matrix of 563 genes in 742 dendritic cells. According to the meta data provided by the original paper, we chose the dendritic cell types as one study target and the two batches of their experiment as another study target. Figure S4(a) is a t-SNE map of original data using Euclidean distance metric. We can see that the original 6 clusters identified in the original paper were well re-discovered. But from the right panel of Figure S4(a), cells from the two batches in each cluster are also separated. This implies that the clustering of the 6 cell types in the original data are the dominate signal in the data, but the batch effect is also obvious. The original paper adopted a step to remove the batch effect. We use the data before removing batch effect to study whether these two orthogonal grouping relations can be revealed when we look at the data from different angles.

**Fig. S4.**
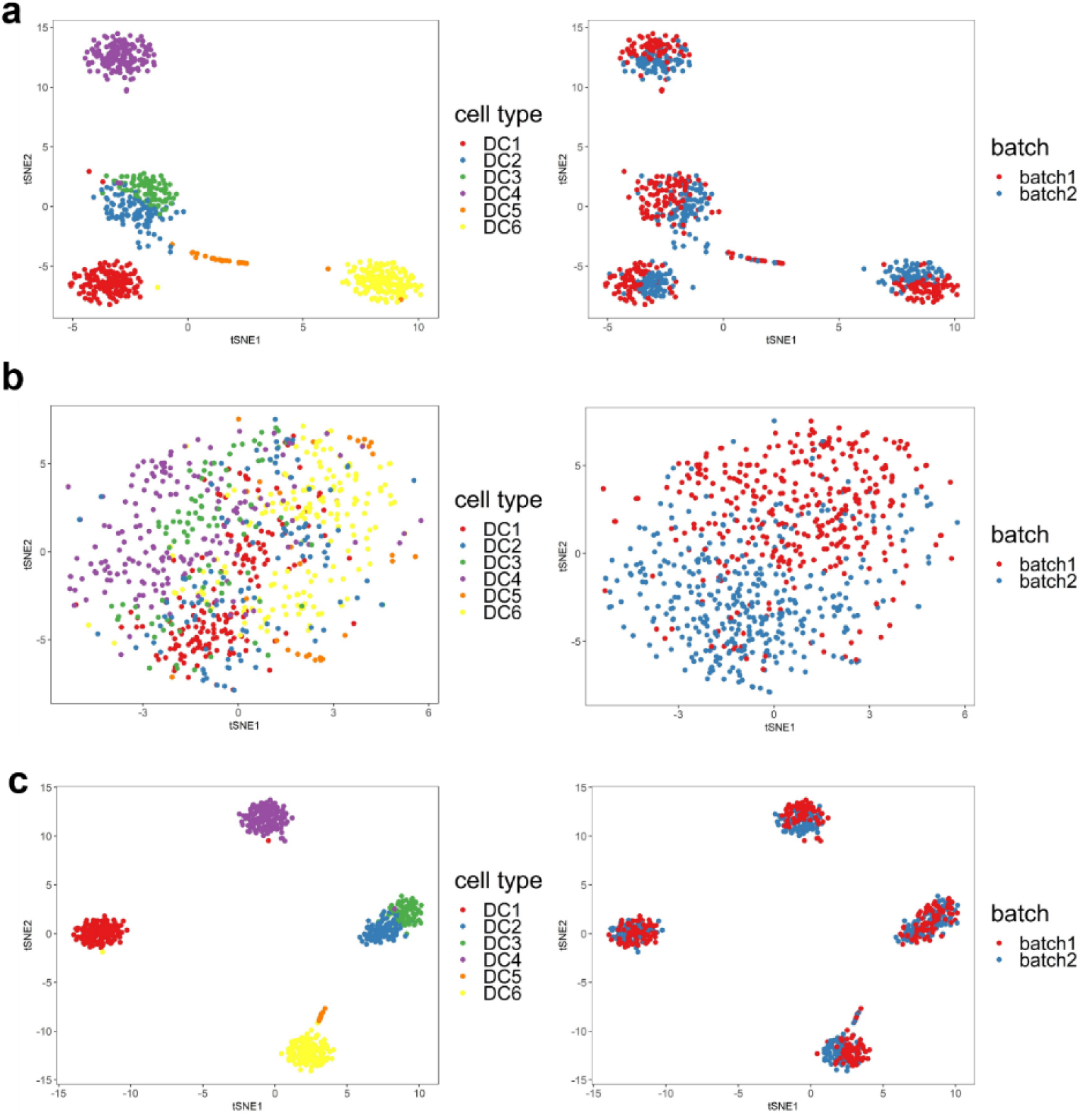
t-SNE maps of dendritic cells under different metrics. (a) t-SNE maps of dendritic cells under Euclidean distance metric. (b) t-SNE maps of dendritic cells under metric learned from batch number label training pairs. (c) t-SNE maps of dendritic cells under metric learned from cell subtype label training pairs.

### Cell batch as similarity criterion

In this dataset, the two batches have almost the same amount of cells, 368 from batch 1 and 374 from batch 2. We select cells randomly to make up training pairs. We select 100 similar training pairs from the same batch and 100 dissimilar pairs across different batches to train new metric. To show how this similarity criterion work on distance metric, we visualize cells via t-SNE respectively based on the Euclidean metric and the learned metric. Fig. S4(b) shows the t-SNE maps under metric learned with the weak supervision of batch signals. We can see that 6 clusters are mixed, but cells of the two batches tend to be distributed in separated regions of the map. The signal of the batches is not as strong as that of the cell types so the batches are not forming isolated clusters even after metric learning.

### Cell subtype as similarity criterion

We then experimented on using the cell type information as weak supervision for scMetric. The numbers of cells in each subtype are very unbalanced as shown in Table S1.

**Table S1.** Numbers of cells in each cell subtype of Dendritic cell dataset

|  |  |
| --- | --- |
| All | 742 |
| DC1(CLEC9A<sup>+</sup>) | 166 |
| DC2(CD1C<sup>+</sup>_A) | 105 |
| DC3(CD1C<sup>+</sup>_B) | 95 |
| DC4(CD1C-CD141-) | 175 |
| DC5(New pop.) | 30 |
| DC6(pDC) | 171 |

Since we don’t have specific domain knowledge on each specific cells, we automatically extracted the weak supervision information in the following manner: We first set a probably of being extracted for each cell so that the total number of representatives in each subtype is roughly the same. This is to guarantee the rare subtypes can also be well represented in the metric learning. With this probably set, we randomly sample cell pairs from all cells. If the two sampled cells are of the same subtype, we put them into the set of similar pairs, and otherwise into the dissimilar pairs. In this way, we collect 6 similar pairs for each cell subtype (36 pairs in total for the 6 subtypes), and 6 dissimilar pairs for each pair of subtypes (180 pairs in total for the 6×5 possible subtype pairs). We arbitrarily pick up 200 pairs of samples from them as the weak supervision data to train scMetric. Figure S4(c) shows the t-SNE map with the learned metric from this weak supervision information. We can see that the clusters become even tighter than those obtained with Euclidean metric, and the batch effect is well removed.

We then checked the top 30 genes that contribute the most in the learned metric. Table S2 shows the list. Among the 242 marker genes for the 6 DC clusters identified in the original paper by requiring AUC>0.85 for discriminating each subtype with the other 5 subtypes, 97 were included in the 563 genes we worked on in this experiment. We found 20 of them were ranked among the top 30 by their importance in the learned metric. The full list of the 563 genes and their importance scores are provided in the supplementary excel file.

**Table S2.**
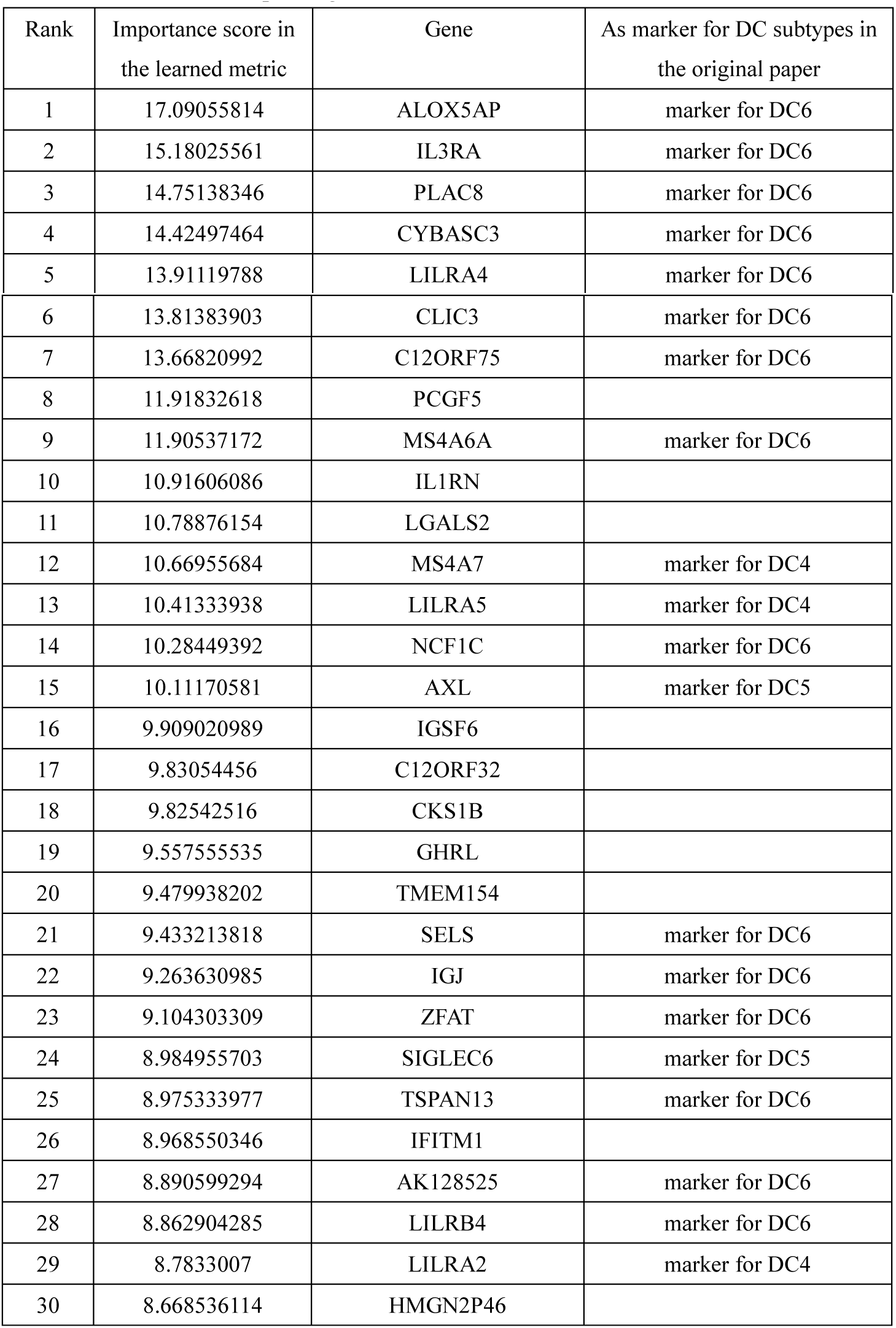
Top 100 genes in the learned metric on the DC data

